# Genomic Background Governs Opposing Responses to Nalidixic Acid Upon Megaplasmid Acquisition in *Pseudomonas*

**DOI:** 10.1101/832428

**Authors:** David A. Baltrus, Caitlin Smith, MacKenzie Derrick, Courtney Leligdon, Zoe Rosenthal, Madison Mollico, Andrew Moore, Meara Clark

## Abstract

Horizontal gene transfer is a significant driver of evolutionary dynamics across microbial populations. Although the benefits of the acquisition of new genetic material are often quite clear, experiments across systems have demonstrated that gene transfer events can cause significant phenotypic changes and entail fitness costs in a way that is dependent on the genomic and environmental context. Here we test for the generality of one previously identified cost, sensitization of cells to the antibiotic nalidixic acid after acquisition of a ∼1Mb megaplasmid, across Pseudomonas strains and species. Overall, we find that the presence of this megaplasmid sensitizes many different Pseudomonas strains to nalidixic acid, but that this same horizontal gene transfer event increases resistance of *Pseudomonas putida* KT2440 to nalidixic acid across assays as well as to ciprofloxacin under competitive conditions. These phenotypic results are not easily explained away as secondary consequences of overall fitness effects and appear to occur independently of another cost associated with this megaplasmid, sensitization to higher temperatures. Lastly, we draw parallels between these reported results and the phenomenon of sign epistasis for *de novo* mutations and explore how context dependence of effects of plasmid acquisition could impact overall evolutionary dynamics and the evolution of antimicrobial resistance.

**Importance:** Numerous studies have demonstrated that gene transfer events (e.g. plasmid acquisition) can entail a variety of costs that arise as byproducts of the incorporation of foreign DNA into established physiological and genetic systems. These costs can be ameliorated through evolutionary time by the occurrence of compensatory mutations, which stabilize presence of a horizontally transferred region within the genome but which also may skew future adaptive possibilities for these lineages. Here we demonstrate another possible outcome, that phenotypic changes arising as a consequence of the same horizontal gene transfer event are costly to some strains but may actually be beneficial in other genomic backgrounds under the right conditions. These results provide new a new viewpoint for considering conditions that promote plasmid maintenance and highlight the influence of genomic and environmental contexts when considering amelioration of fitness costs after HGT events.

## Introduction

Horizontal Gene Transfer (HGT), a catch-all term for the transfer of DNA between individuals, is one of the main drivers of evolutionary dynamics across microbial populations (1– 3). Although there are countless examples where HGT has been shown to increase microbial fitness and/or enable colonization of new niches, it is increasingly appreciated that HGT brings with it a variety of costs that arise as byproducts of the incorporation of foreign DNA into established physiological and genetic systems (4–6). Recent efforts have begun to categorize and characterize phenotypic effects of HGT across systems, and have clearly demonstrated that acquisition of the same genes can lead to different outcomes across genomic backgrounds (7, 8). Such differences in phenotypic outcomes following HGT are analogous if not identical to the concept of sign epistasis in the context of de novo mutations, whereby the fitness effects of a mutation change sign (e.g. from positive to negative) under specific genetic combinations (9– 11). Much like sign epistasis, we currently lack a general understanding of the underlying causes of differences in phenotypic outcomes following HGT, and as a consequence, there is no clear model for predicting if and when acquisition of DNA could be detrimental in some genomic backgrounds but beneficial in others.

We have previously reported numerous phenotypic changes that occur as a consequence of acquisition of a ∼1Mb megplasmid, pMPPla107, by *Pseudomonas* strains (12, 13). In addition to fitness costs correlated with slower growth, acquisition of pMPPla107 alters a variety of other phenotypes in some strains including: increased motility, decreased biofilm formation, decreased thermal tolerance, increased sensitivity to an unidentified compound produced by Pseudomonads, and increased sensitivity to quinolone antibiotics like nalidixic acid and ciprofloxacin. In the particular context of antibiotic sensitivity, it is important to note that current annotations of megaplasmid pMPPla107 contain no known antibiotic resistance genes (14) and that, at present, the mechanistic basis and overall strain specificity of these phenotypic changes remains unknown.

Here, we test for how frequently sensitivity to nalidixic acid is observed across Pseudomonas strains after acquisition of megaplasmid pMPPla107. We show that, although the presence of this megaplasmid increases sensitivity to nalidixic acid in many strain backgrounds, one strain in particular - *Pseudomonas putida* KT2440 − displays increased resistance to nalidixic acid and ciprofloxacin after this HGT event. Therefore, our report provides an additional example whereby the precise phenotypic effects of HGT are dependent on strain background, but also suggests that that antibiotic resistance can be increased by megaplasmid pMPPla107 even in the absence of identifiable resistance genes in the region acquired by HGT. These results, and specifically strain-dependent changes in sign of phenotypic effects of plasmid acquisition, provide a new viewpoint for considering conditions that promote plasmid maintenance and highlight the influence of genomic and environmental contexts when considering amelioration of fitness costs after HGT events.

## Methods

### Bacterial strains and culture conditions

All strains used in this study are listed in Table 1. We note that one of the main *P. stutzeri* strains for previous megaplasmid papers from our group is originally named 28a24 (15), but that in some previous papers from our group we have incorrectly referenced this as strain 23a24 (12, 13). We also note that DBL332 and DBL386 are actually derived from two independent rifampicin resistant mutants selected from strain 28a24 and frozen independently as two different stocks.

**Table 1:**
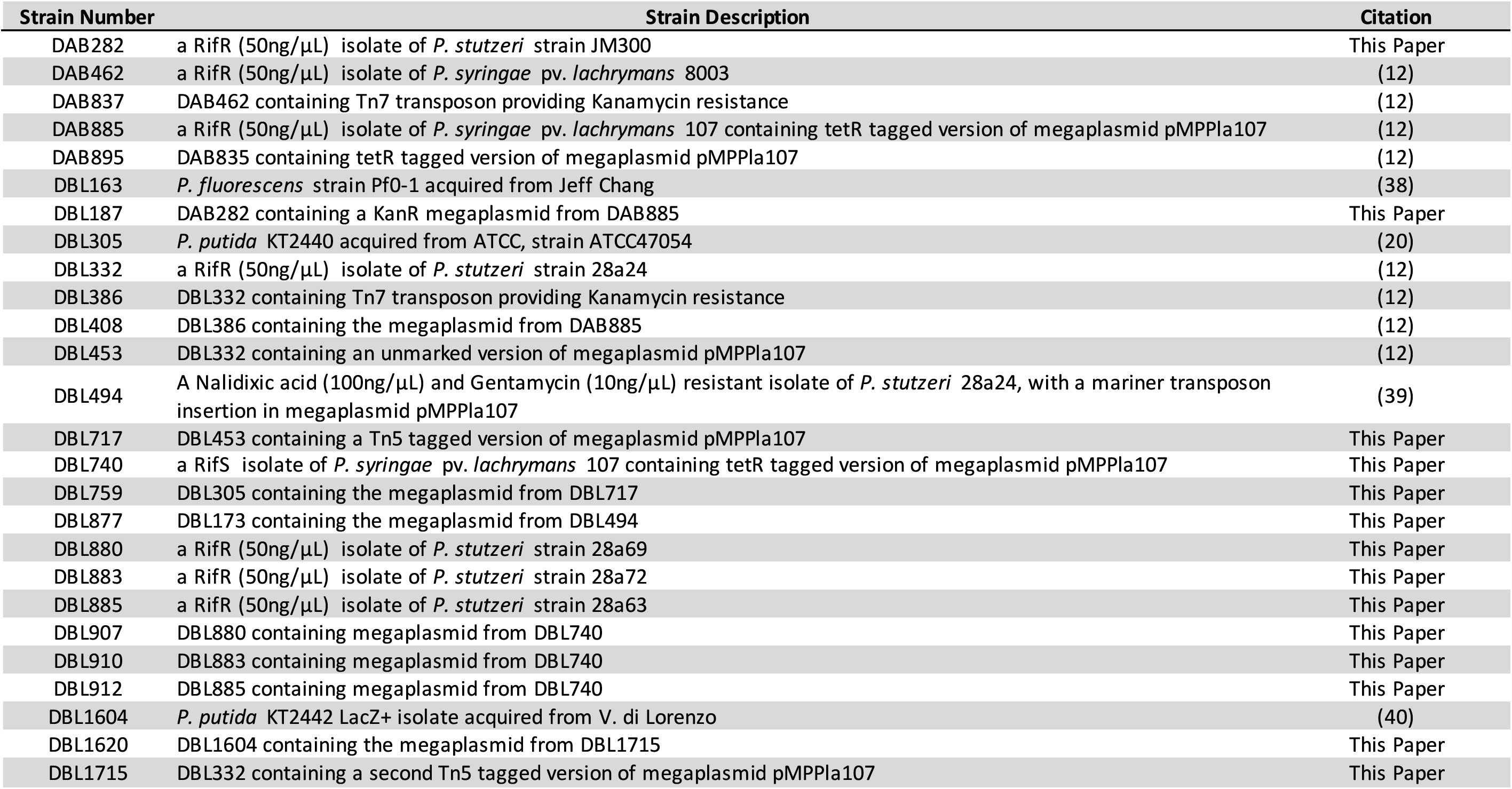
Strains used in this study

Strain propagation of all cultures largely took place at 27°C in King’s B medium supplemented with rifampicin unless otherwise specified. Antibiotics were supplemented into media at the following concentrations when appropriate and unless otherwise specified: 50 µg/mL rifampicin, 10 µg/mL tetracycline, and 20 µg/mL kanamycin. 5-bromo-4-chloro-3-indolyl-β-D-galactopyranoside (Xgal) was supplemented at 40 µg/mL. For competition assays, cultures were supplemented with nalidixic acid at 20µg/mL and 40µg/mL and with ciprofloxacin at 0.5µg/mL. Antibiotic discs used in the diffusion assays contained antibiotics at concentrations of 30 µg/mL for nalidixic acid and 5 µg/mL for ciprofloxacin.

### Kirby-Bauer Disc Diffusion Assays

Strains of interest for each assay were streaked from frozen stocks and incubated on KB media at 27°C for 2 to 3 days. At this point a small clump of cells was picked to liquid KB medium and grown for 24 hours at 27°C with shaking. Cells from each culture were then pelleted in a centrifuge, the supernatant was poured out, and pelleted cells were resuspended in 10Mm MgCl_2_ for measurement of OD_600_. Each culture was then diluted to 0.01 OD_600_ in 10Mm MgCl_2_ and 300uL of each culture was added to 3mL of 0.4% water agar (which was previously melted and allowed to cool until warm to the touch). This bacteria/agar mixture was then poured onto a plain KB agar plate (1.5% agar) and allowed to solidify. Lastly, a disc containing the antibiotic of interest was added to the center of the plate and each was placed face up in an incubator at 27°C. Pairs of megaplasmid +/- strains were always evaluated together within the same assay block.

After 3 days of growth, each plate was taken out of the incubator and scanned. This image was imported into ImageJ (16), and the area of the inhibition zone was measured. An average area was calculated for the megaplasmid-strain within each assay, and individual data points for each pair of megaplasmid + and − strains in each assay were divided by this value in order to normalize the data. Therefore, the average inhibition zone for each megaplasmid-strain within each assay was normalized to a value of 1. Each strain pairing was tested in at least (but usually more than) two independent assays.

### Competitive Fitness Assays

In each assay, *lacZ*+ strains (either DBL1604 or DBL1620, - and + megaplasmid derivatives of strain KT2442, respectively) were competed against the common *lacZ-* strain DBL305 (strain KT2440). Strains of interest for each assay were streaked from frozen stocks for 2 to 3 days (on media containing kanamycin for megaplasmid+ strains) at which point a small clump of cells was picked to liquid KB medium and grown for 24 hours at 27°C with shaking. Cells from each culture were then pelleted in a centrifuge, the supernatant was poured out, and pelleted cells were resuspended in 10Mm MgCl_2_ for measurement of OD_600_. Each culture was then diluted to 1.0 OD_600_ in 10Mm MgCl_2_.

A 10mL master culture was prepared for each set of comparative fitness assays and for each different treatment. To prepare this master culture, 5uL of each 1.0 OD_600_ dilution of the two comparison strains within each assay was added to the 10mL master culture. At this point, 2mL samples of this master culture were added to 4 different test tubes, and the test tubes were placed in a shaking incubator and grown for 24 hours at 27°C. For day 0 measurements, the master mix culture from the control (no antibiotic supplementation) competitions was diluted 1:100 by adding 10µL into 990µL10Mm MgCl2, which was followed by an additional 1:10 dilution. 3 replicate dilution series were made from each master mix culture, and 150µL of each 1:100/1:1000 dilution was plated on LB media containing 40µg/mL Xgal and placed 27°C. After 24 hours of growth each of the 4 replicate cultures was diluted 1:10,000,000 and 1:100,000,000, with 150µL of these dilutions plated on LB media containing 40µg/mL Xgal and placed at 27°C.

White and blue colonies were independently counted for each of the plated day 0 dilutions, these 3 values were averaged together, and the log_10_ value of this average taken as the day 0 colony forming unit count. White and blue colonies were independently counted for each of the day 1 replicate cultures and log_10_ transformed. Relative fitness was calculated as the ratio of the difference between log_10_ transformed white and blue colony counts for each replicate culture at day 1 over the difference of the log_10_ transformed average white and blue colony counts at day 0. Depending on the assay, either two or three assays were carried out with each pair of strains (DBL305/DBL1604 and DBL305/DBL1620), with four replicate cultures per assay.

All competitive fitness assays were carried out in the same manner as described above, with one exception. Since cultures grew more slowly in the presence of 40 µg/mL nalidixic acid, cultures from the nalidixic acid competitions as well as the control competitions were allowed to grow in liquid media at 27°C for 48 hours.

### Statistical Methods

All statistical tests were performed in R version 3.3.0 (17). Scripts for these tests and all underlying data can be found at https:/doi.org/10.5281/zenodo.4409341

## Results

### A Modified Kirby-Bauer Disc Diffusion Assay Replicates Previous Results For Nalidixic Acid Sensitivity Upon Megaplasmid Acquisition

Our previously published experiments thoroughly explored how acquisition of pMPPla107 increases nalidixic acid sensitivity in *Pseudomonas stutzeri* 28a84 (13), and here we demonstrate that this effect is replicated by a simple modified assay based on Kirby-Bauer disc diffusion assays ((18), Figure 1A). Since antibiotics diffuse out from the disc into the media forming a concentration gradient, size of the inhibition halo is positively correlated with sensitivity to nalidixic acid. We screened for nalidixic acid sensitivity in two different backgrounds of *P. stutzeri* strain 28a24, DBL332 and DBL386. Derivative strains of these (DBL453 and DBL408, respectively, which contain megaplasmid pMPPla107) have a larger halo of inhibition around the antibiotic disc containing nalidixic acid than their immediate progenitor strains which lack megaplasmid pMPPla107 (Figure 1B). There is an overall significant effect of megaplasmid on nalidixic acid sensitivity (F_1,36_=132.143, p<0.0001), and there are clear statistical differences between the areas of the inhibition halos for each pair of megaplasmid -/+ strains measured separately (DBL332/DBL453, t-test p<0.001; DBL386/DBL408, t-test p<0.001). Therefore, the acquisition of this megaplasmid sensitizes these strains to this antibiotic.

**Figure 1.**
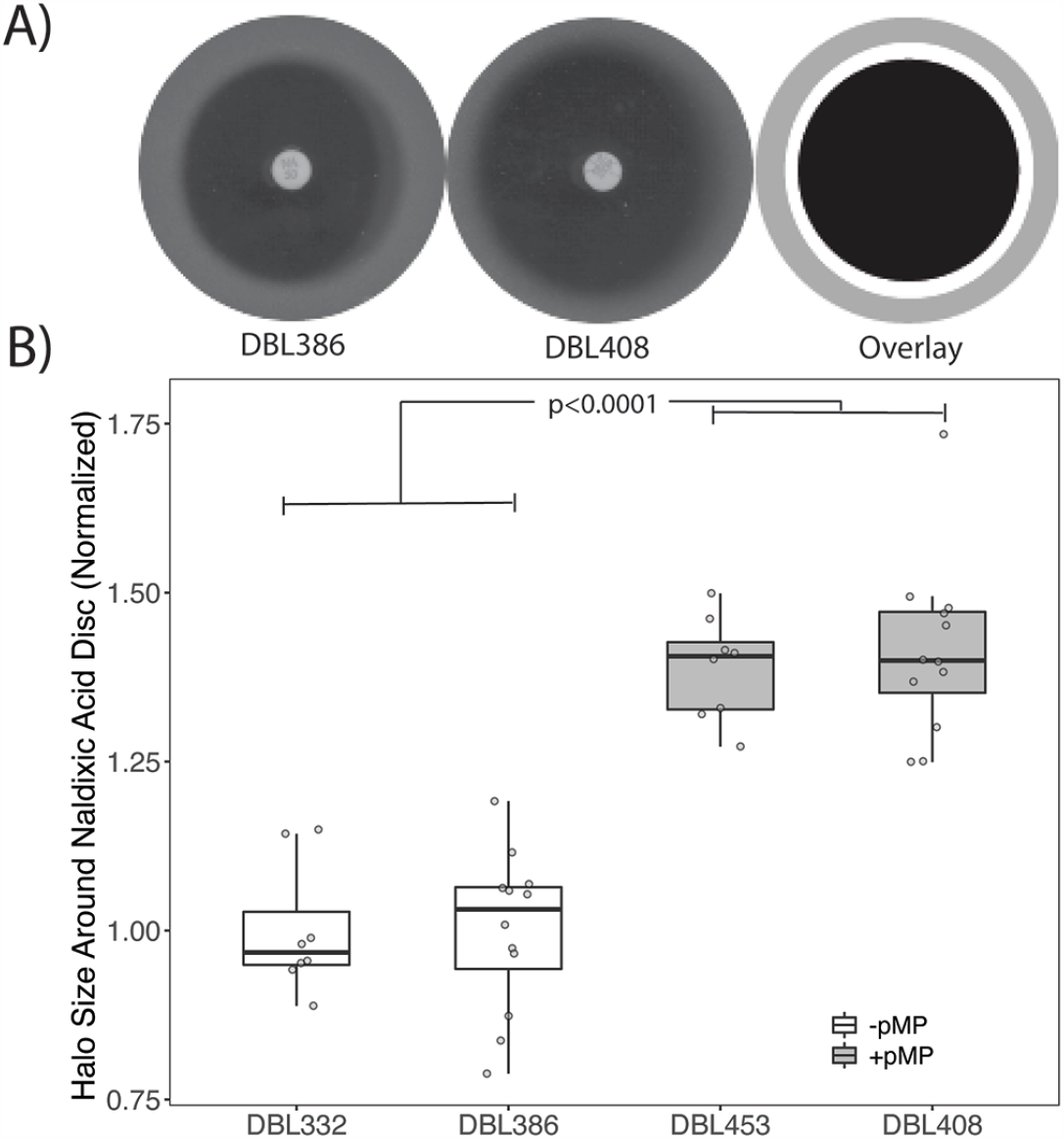
Kirby-Bauer Diffusion Assays Recapitulate A Nalidixic Acid Sensitivity Phenotypes in *Pseudomonas stutzeri*. We have previously demonstrated that acquisition of megaplasmid pMPPla107 increases sensitivity to nalidixic acid in two independent rifampicin resistant derivatives of *P. stutzeri* 28a24, DBL332 and DBL386. Here we show that assays based on diffusion of nalidixic acid out of a disc recapitulate this phenotype. A) Representative scans from a disc diffusion assay from using (from left to right) a strain that either lack (DBL386) or contain (DBL408) megaplasmid pMPPla107. Image areas covered by each of these two representative pictures (grey), the sensitivity halo around the disc containing nalidixic acid and strain DBL386 (black) or strain DBL408 (white) are overlaid onto each other at the far right of this image. The larger the inhibition halo, the greater the sensitivity to nalidixic acid. B) Data from disc diffusion overlay assays comparing *P. stutzeri* strains DBL332 and DBL386 with derivatives which have acquired megaplasmid pMPPla107. Each strain was assayed at least three independent times, with at least 2 (but usually 6) replicates per strain per assay. Halo size was normalized to the relevant “wild type” strain within each assay (DBL332 for DBL453, DBL386 for DBL408) and plotted on the Y-axis. Individual data points are shown for all assays, with box hinges corresponding to first and third quartiles and mean plotted as horizontal lines in the center of the boxes. Strains containing the megaplasmid are plotted in the grey boxes and are more sensitive to inhibition by nalidixic acid (F_1,36_=132.143, p<0.0001) than paired parental strains lacking the megaplasmid (plotted in white). Additionally, there are clear statistical differences between the areas of the inhibition halos for each pair of megaplasmid -/+ strains (DBL332/DBL453, ttest p<.001; DBL386/DBL408, ttest p<0.001)

### Acquisition of pMPPla107 Sensitizes Most, But Not All, Strains to Nalidixic Acid

Extending the Kirby-Bauer diffusion assay design, we tested for sensitivity to nalidixic acid across a variety of other *Pseudomonas stutzeri* strains as well as a *P. syringae* pv. *lachrymans* 8003 and *P. fluorescens* Pf0-1. The *P. syringae* strains are relatively closely related to the strain in which this megaplasmid was originally found, *P. syringae* pv. *lachrymans* 107 (19), and acquisition of the megaplasmid has been previously shown to affect growth of this strain (12). Overall, we find that megaplasmid acquisition increases sensitivity to nalidixic acid across all of these strains as reflected by increased halo size in these Kirby-Bauer assays (Figure 2). However, we also find that the magnitude of this difference appears to be smaller in *P. syringae* than across all of the *P. stutzeri* strains and even smaller for the *P*.*fluorescens* Pf0-1 comparisons. All comparisons between megaplasmid- and megaplasmid+ strains are significantly different (t-test, p<0.001 for each with the exception of p<0.05 for the Pf0-1 comparison). We have also included representative scans from each of these comparisons in Supplemental Figure 1 to demonstrate relative levels of antibiotic sensitivity (found at https://doi.org/10.6084/m9.figshare.10257704.v2)

**Figure 2.**
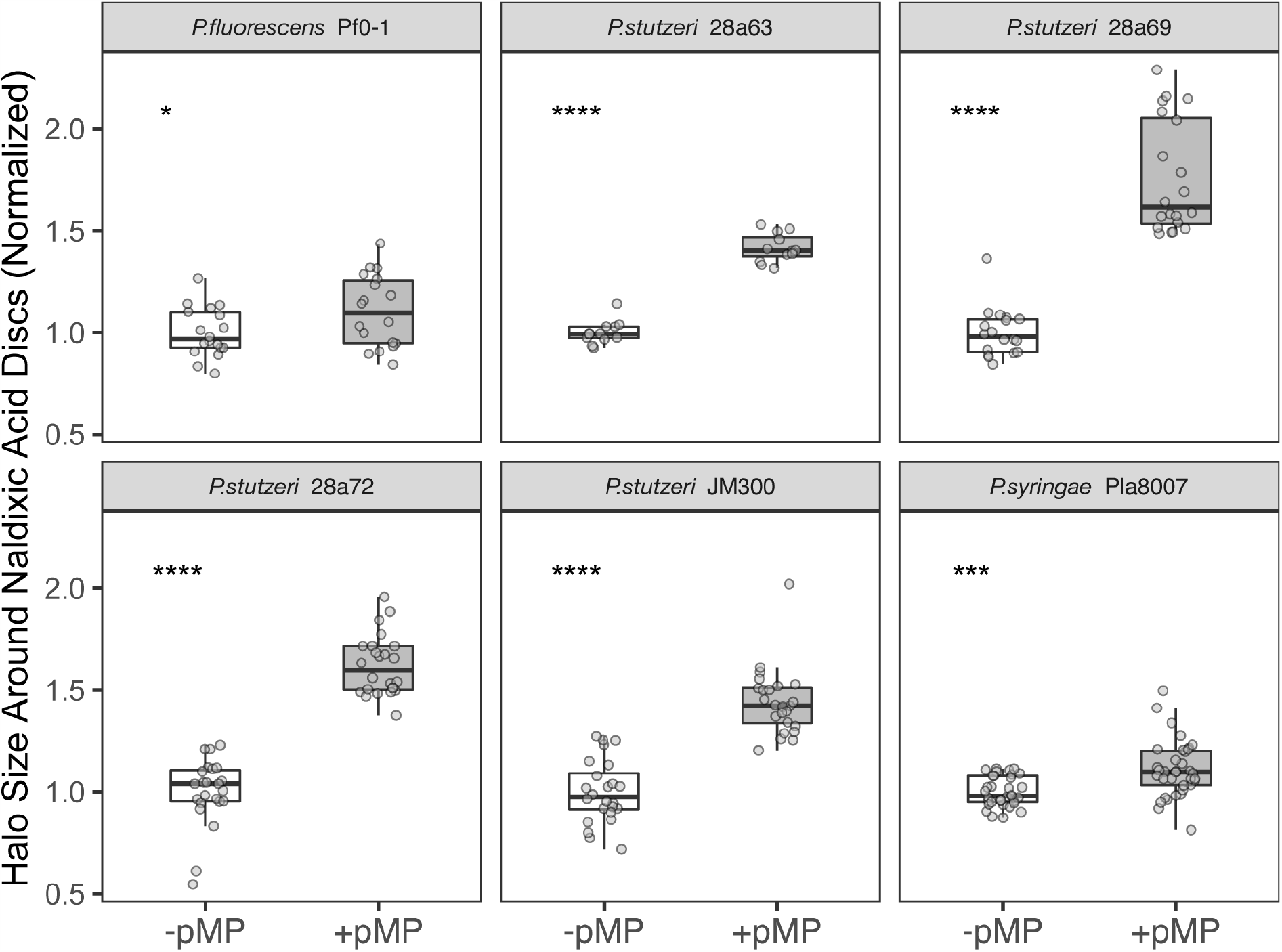
Acquisition of Megaplasmid pMPPla107 Sensitizes Numerous Pseudomonas strains to Nalidixic Acid. Data is shown for disc diffusion overlay assays comparing *P. stutzeri* strains DBL880, DAB282, DBL883, and DBL885 with derivatives which have acquired megaplasmid pMPPla107 (DBL907, DBL287, DBL910, DBL912 respectively). Data is also shown for derivatives of *P. syringae* pv. *lachrymans* strain 8003 and *P. fluorescens* Pf0-1 which either lack (DBL462 and DBL163, respectively) or contain (DBL895 and DBL877, respectively) megaplasmid pMPPla107. Strain names of the original isolates of genomic backgrounds lacking the megaplasmid are shown in grey headers. Data for “wt” strains are plotted in white boxes, while data for strains containing the megaplasmid are plotted in grey boxes. Each strain was assayed at least three independent times, with at least 4 (but usually 6) replicates per strain per assay. Two additional assays were performed for *P. syringae* strains. Halo size was normalized to the relevant “wild type” strain within each assay and plotted on the Y-axis. Individual data points are shown for all assays, with box hinges corresponding to first and third quartiles and mean plotted as horizontal lines in the center of the boxes. In each case, presence of megaplasmid pMPPla107 sensitizes strains to nalidixic acid as shown by increased halos of inhibition. Asterisks indicate statistical significance by t-test: >0.05=ns, *=<0.05, **=<0.01, ***=<0.001, ****=<0.0001. All intrastrain comparisons between parent strains that lack the megaplasmid and derivative strains containing the megaplasmid are significantly different (p<0.001) by t-test with the exception of the Pf0-1 comparison which was p<0.05 by t-test.

### Acquisition of Megaplasmid pMPPla107 by *P. putida* Increases Resistance to Nalidixic Acid

Using the same diffusion assays as above, we find that acquisition of megaplasmid pMPPla107 by *Pseudomonas putida* KT2440 repeatedly increased resistance of this strain to nalidixic acid (as reflected by a smaller halo in megaplasmid+ strains in Figure 3). This effect was replicated in an independently created pair of *P. putida* strains originally derived from *P. putida* KT2442, where megaplasmid pMPPla107 was tagged with a different antibiotic resistance marker (Figure 3A). *P. putida* KT2442 is a spontaneous rifampicin resistant isolate of strain KT2440 (20). Across both strain pairs, presence of megaplasmid pMPPla107 increases resistance of *P. putida* strains to nalidixic acid (F_1,85_=147.57, p<0.0001) and there are clear statistical differences between the areas of the inhibition halos for each pair of megaplasmid -/+ strains (DBL305/DBL759, t-test p<.001; DBL1604/DBL1620, t-test p<0.001). However the magnitude of this effect does appear to be somewhat strain specific (Figure 3A, F_2,83_=58.36, p<0.001)).

**Figure 3.**
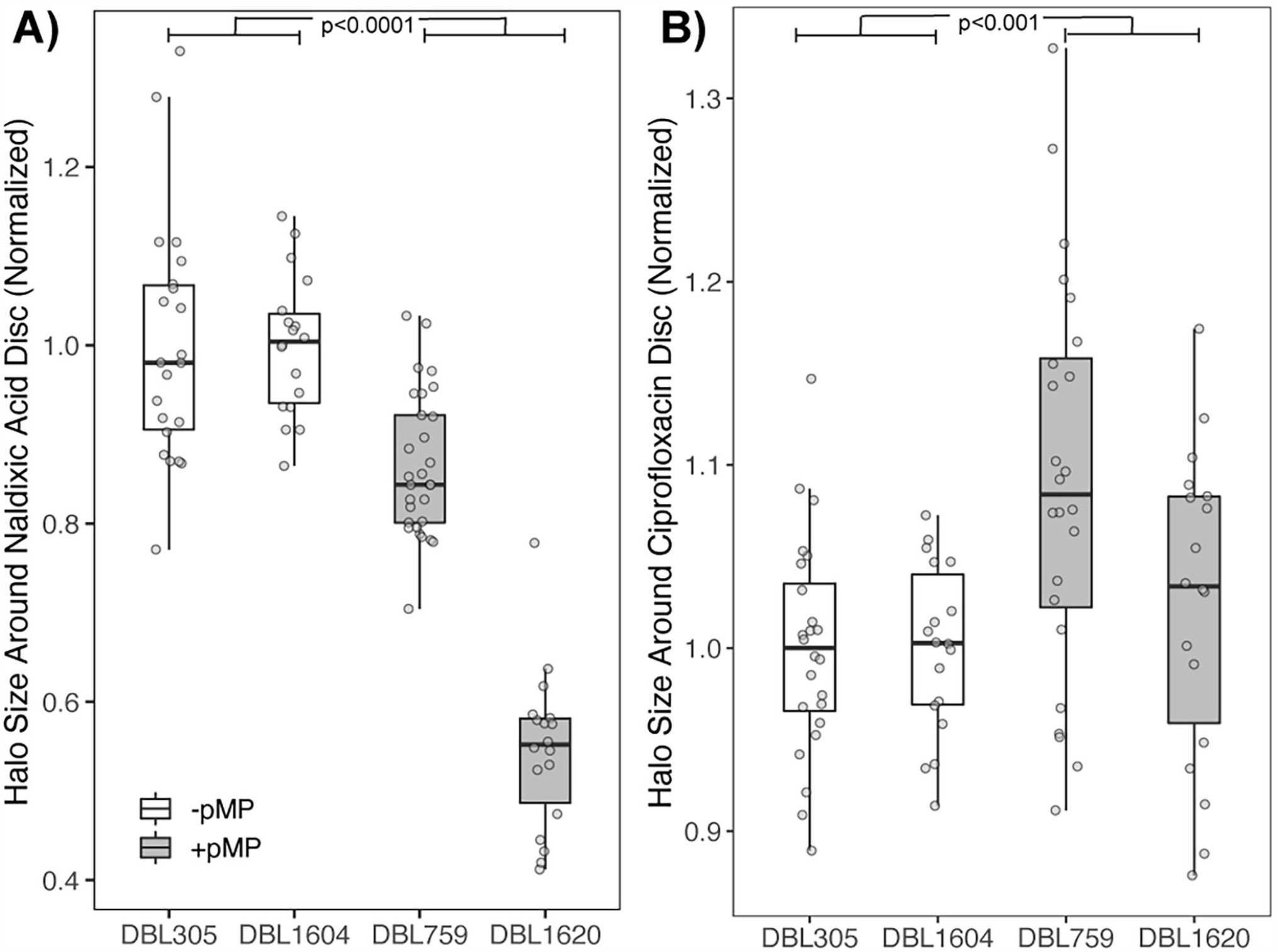
Acquisition of Megaplasmid pMPPla107 by *P. putida* Increases Resistance to Nalidixic Acid but Not Ciprofloxacin in Disc Diffusion Assays. Data is shown for disc diffusion overlay assays comparing *P. putida* strains DBL305 (KT2440) and DBL1604 (KT2442) with derivatives which have acquired megaplasmid pMPPla107 (DBL759 and DBL1620 respectively). Each strain was assayed at least three independent times for each antibiotic, with 6 replicates per strain per assay. Halo size was normalized to the relevant “wild type” strain within each assay and plotted on the Y-axis. Individual data points are shown for all assays, with box hinges corresponding to first and third quartiles and mean plotted as horizontal lines in the center of the boxes. Data for “wt” strains are plotted in white boxes, while data for strains containing the megaplasmid are plotted in grey boxes. A) Presence of megaplasmid pMPPla107 increases resistance of *P. putida* strains to nalidixic acid as shown by decreased halos of inhibition (F_1,85_=147.57, p<0.0001). Moreover, there are clear statistical differences between the areas of the inhibition halos for each pair of megaplasmid -/+ strains (DBL305/DBL759, t-test p<.001; DBL1604/DBL1620, t-test p<0.001) B) Presence of megaplasmid pMPPla107 appears to sensitize *P. putida* strains to ciprofloxacin as shown by increased halos of inhibition (F_1,80_=12.99, p<0.001). However, although ciprofloxacin halo inhibition areas statistically differ between strains DBL305 and DBL759 (ttest, p<0.001), there is no clear difference in the inhibition halos between strains DBL1604 and DBL1620 (ttest, p=0.29).

Nalidixic acid and ciprofloxacin are both quinolone antibiotics that have the same mechanism of action and cause bacterial cell death by inhibiting the ability of gyrase or topoisomerase IV to covalently connect DNA strands after unwinding. Using the same Kirby-Bauer diffusion assays as above, but instead replacing nalidixic acid discs with ciprofloxacin discs, we find that acquisition of megaplasmid pMPPla107 does not clearly increase resistance to this antibiotic (Figure 3B). Although the results are subtle, strains appear to be overall sensitized by the megaplasmid to ciprofloxacin (F_1,80_=12.99, p<0.001). However, only one of the megaplasmid+/- strain pairs is significantly different when evaluated individually (DBL305/DBL759 t-test, p<0.001; DBL1604/DBL1620 t-test, p=0.29).

### A General Cost of Acquisition of Megaplasmid pMPPla107 by *P. putida*

We have previously shown that acquisition of pMPPla107 by *P. stutzeri* 28a84 leads to dramatic fitness costs when measured by growth rate and by competitive fitness assay, and have demonstrated that growth in the presence of nalidixic acid increases this fitness cost (12, 13). We therefore carried out competitive fitness assays of differentially marked *P. putida* strains in order to test whether megaplasmid acquisition also affected competitive growth in this strain background. As one can see in Fig. 4A, there is no measurable difference in competitive fitness between the *lacZ+* background (KT2442) and the wild type background (KT2440) (t-test compared to value of 1, p=0.14). However, in media without any antibiotic supplementation, megaplasmid acquisition leads to a ∼10% loss in competitive fitness over one growth cycle (Figure 4A), and this difference is significantly different from the control comparison (F_1,20_=600.3 p<0.0001). Therefore, increased nalidixic acid resistance as seen in the assays above occurs even though megaplasmid acquisition significantly lowers strain growth rates in the absence of antibiotics.

**Figure 4.**
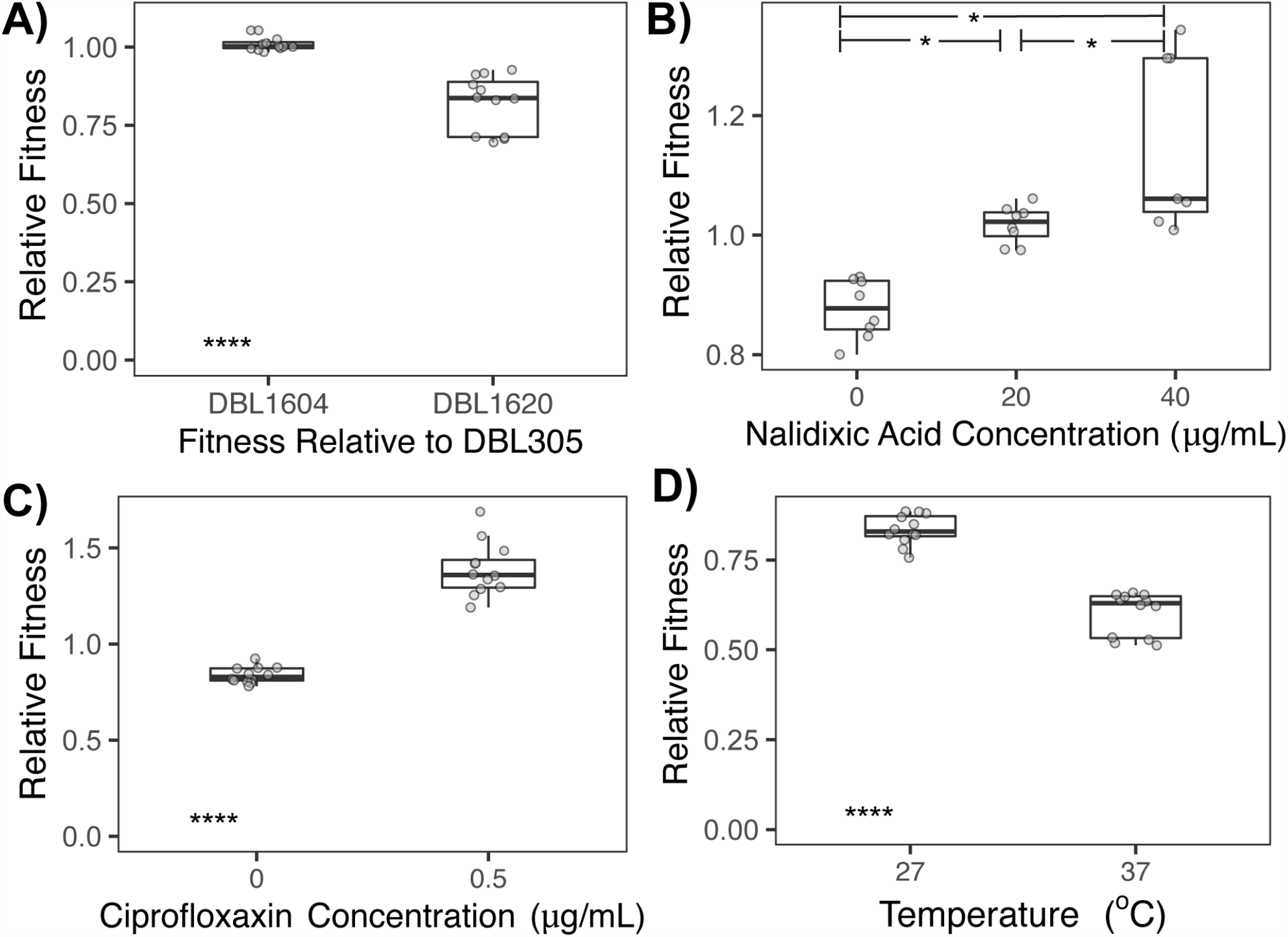
Contrasting Effects of Acquisition of Megaplasmid pMPPla107 in *P. putida* Across Different Environments as Measured by Competitive Fitness Assays. Competitive fitness assays in liquid KB media were carried out between strain DBL305 (*P. putida* KT2440) and *lacZ*+ strains that either lack (DBL1604) or contain (DBL1620) megaplasmid pMPPla107. At least two competitive fitness assays were carried out for each pairing with 4 replicates per assay in all but one case (one assay at Nal40 only had 3 replicates). Individual data points are shown for all assays, with box hinges corresponding to first and third quartiles and mean plotted as horizontal lines in the center of the boxes. Asterisks indicate statistical significance by t-test: >0.05=ns, *=<0.05, **=<0.01, ***=<0.001, ****=<0.0001. A) Although there is no measurable difference in competitive fitness due to the *lacZ+* marker (assays between DBL305 and DBL1604), presence of the megaplasmid leads to a ∼10% fitness cost during growth in KB medium (F_1,20_=600.3 p<0.0001). B) Although the ∼10% fitness cost of the megaplasmid is again apparent during competition between DBL305 and DBL1620 in unsupplemented KB medium, strain 1620 (containing megaplasmid pMPPla107) displays increasingly higher relative fitness compared to strain DBL305 when media is supplemented with nalidixic acid at either 20 or 40 ng/uL ((F_2,17_=264.82 p<0.0001), all comparisons significant at p<0.05 by Tukey’s HSD) or with C) ciprofloxacin at 0.5µg/mL (F_1,18_=178.89 p<0.0001). D) The fitness cost of megaplasmid carriage greatly increases, to >50%, when competitions are carried out at higher temperatures of 37°C (F_1,18_=319.38 p<0.0001).

### Supplementation with either Nalidixic Acid or Ciprofloxacin Shifts The Results of Competitive Fitness Assays In *P. putida*

Since megaplasmid acquisition by *P. putida* appears to increase nalidixic acid resistance according to Kirby-Bauer diffusion assays, we tested whether supplementation of media with nalidixic acid could shift the outcomes of competitive fitness assays between wild type and a megaplasmid containing strain. In the absence of any antibiotic supplementation, we are able to independently recapitulate the fitness costs of megaplasmid acquisition reported above, and again find a significant cost of megaplasmid acquisition when measured by competitive fitness (Figure 4B). However, fitness relationships between strain pairs that lack or contain the megaplasmid shift when nalidixic acid is supplemented into media (F_2,17_=264.82 p<0.0001). At 20µg/mL, a ∼10% fitness cost of the megaplasmid changes to a slight benefit for carriage of the plasmid. Moreover, we see increasing fitness according to these competitive assays for the megaplasmid strain at even higher levels of nalidixic acid (40µg/mL), although the exact magnitude of this effect appears to be quite variable (Figure 4B). Competitive fitness measurements are significantly different between each concentration of antibiotics (Tukey’sHSD, p<0.05). Therefore, megaplasmid acquisition by *P. putida* significantly increases resistance to nalidixic acid as measured by both Kirby-Bauer diffusion assays as well as by competitive fitness assays. Despite somewhat somewhat confounding results from disc diffusion assays from these same strains with ciprofloxacin, but consistent with the results of competitive assays supplemented with nalidixic acid, we also find that acquisition of megaplasmid pMPPla107 by *P. putida* increases the competitive ability of this strain background in the presence of 0.5ng/uL ciprofloxacin (Figure 4C; F_1,18_=178.89, p<0.0001).

### Megaplasmid Dependent Resistance to Nalidixic Acid in P. putida Is Not Positively Correlated with Temperature Sensitivity

Acquisition of megaplasmid pMPPla107 by *P. stutzeri* also renders this strain more sensitive to relatively higher temperatures (35-37°C, (12)). To test for this effect in *P. putida*, we carried out competitive fitness assays as above at both 27 and 37°C. Much like our previous results in *P. stuzeri*, the cost of megaplasmid acquisition in *P. putida* is greater when strains undergo competitive fitness assays at 37°C compared to 27°C (F_1,18_=319.38 p<0.0001).

## Discussion

There is a growing interest in predicting the evolutionary dynamics within microbial populations, as a means to drive evolution within populations and communities towards beneficial outcomes as well as a way to control or limit change within extant populations (21). Horizontal Gene Transfer (HGT) is well recognized as a powerful driver of evolutionary dynamics within microbial communities (1–3), and as such, should be incorporated into scenarios for predicting future evolutionary dynamics. Plasmid conjugation and acquisition is a major contributor to HGT in bacterial populations and communities, and plasmids are well known to provide large-scale evolutionary benefits like enabling antibiotic resistance (2, 3, 22). However, plasmid carriage can also entail a variety of fitness costs, which can be manifest as lowered fitness or other phenotypic changes for plasmid bearing strains, in specific environments (4–6). The presence of costs associated with plasmid carriage has led to speculation about the conditions that enable plasmid maintenance within populations through time, with one likely scenario being that costs associated with plasmid carriage are ameliorated over time through subsequent and potentially rapid compensatory adaptation (23, 24). Although not all HGT events entail fitness costs, rapid amelioration of fitness costs associated with plasmid acquisition could lead to genetic and phenotypic tradeoffs that fundamentally shift future evolutionary trajectories for the affected populations. As such, and with a lofty goal of ultimately being able to predict evolution across bacterial populations, it is therefore critical to understand how costs of plasmid carriage are affected by genetic, genomic, and environmental contexts.

Here we demonstrate that the direction of one previously observed phenotype associated with acquisition of megaplasmid pMPPla107, sensitivity to nalidixic acid in *Pseudomonas (13)*, depends upon the interactions between this megaplasmid and genomic background of the strain of interest. Across almost all surveyed strains, acquisition of pMPPla107 sensitizes cells to nalidixic acid, but in *P. putida* KT2440 this same horizontal transfer event increases resistance to this antibiotic. These contrasting phenotypic effects occur in the absence of identifiable megaplasmid genes which could be hypothesized to influence antibiotic sensitivity and resistance. Thus, this is not simply a scenario where a common antibiotic resistance gene found on the megaplasmid is active in one background compared to the others, but rather a situation where the phenotypic effects of genes present on a megaplasmid are polar opposites under certain environments in ways that depend on the chromosomal background. Also important, these same phenotypic benefits are seen when megaplasmid + and - strains of *P. putida* are competed together in fitness assays. We also demonstrate that these same competitive fitness effects can be seen when megaplasmid + and - strains of *P. putida* are competed in the presence of ciprofloxaxin, an antibiotic which has the same mechanism of action as nalidixic acid. These results suggest that the mechanism underlying changes in sensitivity to nalidixic acid and/or ciprofloxacin could be due to alterations in the activity or amounts of gyrase and topoisomerase activity within *P. putida* cells after acquisition of an additional 1Mb of DNA.

Although we did not see a clear and similar benefit to megaplasmid acquisition by *P. putida* using disc assays for ciprofloxacin, it is possible that discrepancies could arise because of differences in the concentration of ciprofloxacin across both types of assay. In disc assays, ciprofloxacin diffuses out into the plate from a disc seeded with 5µg/mL of this antibiotic whereas competitive fitness assays took place at 0.5ng/mL concentrations. We also observed that there was a sharp gradient at ciprofloxacin concentrations in competitive fitness assays whereby above 0.5ng/mL all strains were inhibited in growth (data not shown). It is possible that there is a small window to dial in the benefits of megaplasmid pMPPla107 across concentrations of ciprofloxacin and that the concentrations in the disc assay do not enable the resolution to pick apart clear differences in strains. It is also possible that the benefits of acquisition of pMPPla107 for ciprofloxacin resistance by *P. putida* are subtly dependent on the precise type of assay and potentially on the exact strain background being assayed. For instance, only one of the two strain pairings used for *P. putida* (DBL305 and DBL759) was significantly different in the ciprofloxacin disc assays and it’s possible that secondary mutations that occurred during construction of these strains are responsible for this result. Ultimately, and regardless of confounding results between multiple types of assays for ciprofloxacin, the consistency of the nalidixic acid results across assays and support from competitive fitness assays, clearly demonstrate that the phenotypic consequences of acquisition of pMPPla107 by *P. putida* are fundamentally different than that in *P. stutzeri*.

Quinolone antibiotics have been used as a means of curing a variety of plasmids from many different bacterial strains, given that they inhibit the functions of topoisomerase IV as well as gyrase (25), which implies that plasmid driven sensitivity to quinolones could be a widespread phenomenon (26). These previously reported effects are thought to be mediated by stress at the level of gyrase and topoisomerase function and should therefore likely be affected by either nalidixic acid or ciprofloxacin (26). At present there are no clear mechanistic explanations for differing phenotypic responses of *Pseudomonas* strains to acquisition of pMPPla107 and any potential scenario must involve interactions between genes on both the chromosome and megaplasmid with one outcome of this interaction being differential sensitivity to nalidixic acid. While annotations of pMPPla107 do not point to any genes directly implicated in nalidixic acid resistance (14), this megaplasmid does contain two genes that together are annotated as *parC* and *parE* (locus tags PLA107_031430 and PLA107_031415 encoding topoisomerase IV subunits A and B, respectively). Interestingly, blastP comparisons of all complete Pseudomonas genomes through Pseudomonas.com (27), suggests that the megaplasmid *parC* is closely related to chromosomal orthologs from *P. putida* (∼75% amino acid similarity) while *parE* is closely related to chromosomal orthologs in *P. stutzeri* (∼82% amino acid similarity). Moreover, quinolones are known to bind to topoisomerase subunit A to disrupt its activity with resistance mutations often occurring in this subunit across different bacteria (25). It is certainly possible that negative and positive interactions between megaplasmid encoded topoisomerase IV subunits with their chromosomal orthologs in each strain background could mediate the differences reported here in response to quinolone antibiotics and this will be a focus of future studies.

Gene annotations also suggest that the megaplasmid contains multiple efflux pumps. While the substrates for these pumps are currently unknown, it is plausible that one could specifically transport quinolone antibiotics outside of the cell in *P. putida*. Indeed, there is precedence for efflux pumps in *Pseudomonas aeruginosa* and other bacteria to differentially transport quinolone antibiotics (28, 29). Following this line of thought, it would then be possible that either the particular regulation or activity of this pump differs based on chromosomal background. There may also be a mechanism to differentially detoxify nalidixic acid and ciprofloxacin that resides on the chromosome (whether through efflux pumps or other means), but where the activity of which is differentially affected by megaplasmid encoded pathways. Perhaps there is a global regulator which is similarly affected by megaplasmid presence, but where the downstream phenotypic effects differ across strain backgrounds. For example, megaplasmid presence could uniformly lead to upregulation of AmrZ across Pseudomonads, but upregulation of AmrZ may trigger opposing phenotypic effects in the context of quinolone resistance in different strain backgrounds (30). It is also possible that phenotypic outcomes of increased sensitivity and resistance to quinolone in the presence of pMPPla107 are due to mechanistically different effects. Under this case, the default condition might be that plasmid acquisition causes cells to become more sensitive to certain antibiotics, but that in some backgrounds (in this case *P. putida*) there is an additional pathway or efflux pump that is differentially affected by megaplasmid presence and which enables cells to overcome sensitization found in other *Pseudomonas* species.

As we have demonstrated previously in *P. stutzeri*, we show that acquisition of megaplasmid pMPPla107 is associated with a large (∼10%) fitness cost in *P. putida* in conditions without antibiotic supplementation. Therefore, differing responses to nalidixic acid cannot simply be explained as a secondary consequence of overall growth defects due to acquisition of pMPPla107 and there must be some other inherent systems level effect that creates opposing phenotypic responses across strain backgrounds upon acquisition of this megaplasmid. We further demonstrate that acquisition of pMPPla107 by *P. putida* also increases sensitivity of this strain background to relatively higher temperatures. These contrasting results of megaplasmid acquisition on fitness across multiple environments in *P. putida* suggest that the multiple previously observed fitness costs associated with megaplasmid pMPPla107 (sensitization to antibiotics and higher temperatures) have independent underlying mechanistic bases, at least in *P. putida*.

Specific evolutionary responses to selective pressures are driven by the availability of adaptive mutations within the context of an adaptive landscape (31). Costs associated with plasmid acquisition can rapidly be ameliorated by new mutations that occur on either the plasmid or throughout the rest of the genome, with the speed of compensation correlated with selection pressures acting on the plasmid bearing population (23, 24). Extrapolation from work focused on the molecular and theoretical basis of adaptation predicts that the target size or number of potential compensatory mutations should increase with the cost of plasmid carriage (32). Given that results presented here as well as those reported in other recent papers strongly suggest that costs of plasmid carriage change depending on the environment (33), it is therefore likely that both the speed and molecular basis of amelioration of plasmid costs will greatly differ across environments. Likewise, since the costs and phenotypic consequences of plasmid carriage differ dramatically across genomic backgrounds, there will likely be substantial variation in both the evolutionary paths and molecular basis of compensatory mutations for the exact same plasmid within different strains. The phenotypic effects reported here suggest that plasmids could be differentially maintained or lost across bacterial populations in ways that depend on genotype by environment interactions rather than on simply based on the presence or absence of beneficial genes. Results presented here therefore also strongly bear on our understanding and predicting the evolutionary forces that enable plasmid sharing across microbial communities.

The phenomenon reported here suggests that plasmid acquisition could fundamentally change evolutionary dynamics of antibiotic resistance in a strain dependent way. For instance, there will likely be a different suite of resistance mutations available towards nalidixic acid in *P. putida* KT2440 depending on whether this genomic background contains pMPPla107 or not. This is very much in the vein of sign epistasis when discussing phenotypic effects of *de novo* mutation and evolutionary pathways of adaptation (9, 10, 34, 35). It is also currently unknown how phenotypic effects of particular resistance mutations are altered in the presence of this megaplasmid, but we note that the values evaluated here (20-40ug/mL for Nalidixic acid and 0.5 ug/mL for ciprofloxacin) are relatively close to relevant minimum inhibitory concentrations of hospital isolates of *P. putida* (36). Given that there will likely be tradeoffs in gyrase activity with mutations towards quinolone resistance, one could envision scenarios where acquisition of this megaplasmid simply adds to the level of resistance to nalidixic acid, has no phenotypic effect in a background that already contains nalidixic acid resistance mutations, or even sensitizes cells to nalidixic acid. For example, if a mutation arises in *P. putida* KT2440 that provides resistance to nalidixic acid at 100ng/uL, it is possible that further acquisition of pMPPla107 may either provide no benefit (Minimum Inhibitory Concentration (MIC)+pMPPla107=100ng/uL), an additive increase to resistance (MIC+pMPPla107>100ng/uL), or sensitizes this strain (MIC+pMPPla107<100ng/uL). Future experiments will be necessary to differentiate between these possibilities.

In conclusion, we report here that acquisition of megaplasmid pMPPla107 leads to opposing phenotypic effects of strains in the presence of nalidixic acid but not ciprofloxacin. In most *Pseudomonas* strains evaluated here (which represent a relatively diverse slice from the broader *Pseudomonas* phylogeny (37)), there is a cost to acquisition of pMPPla107 in that cells containing this megaplasmid are sensitized to this antibiotic. However, at least in *P. putida*, acquisition of pMPPla107 increases resistance to nalidixic acid in the absence of identifiable genes involved in antimicrobial resistance. We highlight that these results of HGT are analogous to the phenomenon of sign epistasis when discussing phenotypic effects of *de novo* mutations within a genome. Therefore, as with sign epistasis, it is therefore likely that some HGT events will lead to fundamental shifts in the adaptive landscape in ways that depend on both the genomic background and environment. Depending on how widespread such results are, context dependence of phenotypic effects will potentially increase uncertainty in models and predictions for the evolutionary dynamics of microbial adaptation after horizontal gene transfer events as well as antimicrobial resistance.

## Data Availability

Scripts for recreating all figures and statistical tests using R, as well as all underlying data for these scripts, can be found at **https:/doi.org/10.5281/zenodo.4409341**. Additionally, supplemental Figure 1 can be found at https://doi.org/10.6084/m9.figshare.10257704.v2.

## Acknowledgements

The authors would like to thank Kevin Dougherty and Dr. Brian Smith for help with conceptualization and and implementation of experiments.

## Notes

https://doi.org/10.6084/m9.figshare.10257704.v1

### Competing Interest Statement

The authors have declared no competing interest.

### Summary of Updates

Small typos fixed and table included.

## References

1. Soucy SM, Huang J, Gogarten JP. 2015. Horizontal gene transfer: building the web of life. Nat Rev Genet 16:472–482.

2. Hall JPJ, Brockhurst MA, Harrison E. 2017. Sampling the mobile gene pool: innovation via horizontal gene transfer in bacteria. Philos Trans R Soc Lond B Biol Sci 372.

3. Brockhurst MA, Harrison E, Hall JPJ, Richards T, McNally A, MacLean C. 2019. The Ecology and Evolution of Pangenomes. Current Biology.

4. Baltrus DA. 2013. Exploring the costs of horizontal gene transfer. Trends Ecol Evol 28:489– 495.

5. Carroll AC, Wong A. 2018. Plasmid persistence: costs, benefits, and the plasmid paradox. Can J Microbiol 64:293–304.

6. San Millan A, MacLean RC. 2017. Fitness Costs of Plasmids: a Limit to Plasmid Transmission. Microbiol Spectr 5.

7. Kottara A, Hall JPJ, Harrison E, Brockhurst MA. 2018. Variable plasmid fitness effects and mobile genetic element dynamics across Pseudomonas species. FEMS Microbiol Ecol 94.

8. Alonso-del Valle A, León-Sampedro R. 2020. The distribution of plasmid fitness effects explains plasmid persistence in bacterial communities. bioRxiv.

9. Silva RF, Mendonça SCM, Carvalho LM, Reis AM, Gordo I, Trindade S, Dionisio F. 2011. Pervasive sign epistasis between conjugative plasmids and drug-resistance chromosomal mutations. PLoS Genet 7:e1002181.

10. Weinreich DM, Watson RA, Chao L. 2005. Perspective: Sign epistasis and genetic constraint on evolutionary trajectories. Evolution 59:1165–1174.

11. JAGM de Visser, Cooper TF, Elena SF. 2011. The causes of epistasis. Proc Biol Sci 278:3617–3624.

12. Romanchuk A, Jones CD, Karkare K, Moore A, Smith BA, Jones C, Dougherty K, Baltrus DA. 2014. Bigger is not always better: transmission and fitness burden of 1 MB Pseudomonas syringae megaplasmid pMPPla107. Plasmid 73:16–25.

13. Dougherty K, Smith BA, Moore AF, Maitland S, Fanger C, Murillo R, Baltrus DA. 2014. Multiple Phenotypic Changes Associated with Large-Scale Horizontal Gene Transfer. PLoS ONE.

14. Smith BA, Leligdon C, Baltrus D. 2018. Just the Two of Us? A Family of Pseudomonas Megaplasmids Offers a Rare Glimpse Into the Evolution of Large Mobile Elements. bioRxiv.

15. Sikorski J, Teschner N, Wackernagel W. 2002. Highly different levels of natural transformation are associated with genomic subgroups within a local population of Pseudomonas stutzeri from soil. Appl Environ Microbiol 68:865–873.

16. Schneider CA, Rasband WS, Eliceiri KW. 2012. NIH Image to ImageJ: 25 years of image analysis. Nat Methods 9:671–675.

17. R Core Team. 2016. R: A language and environment for statistical. R Foundation for Statistical Computing, Viena, Austria, Vienna, Austria.

18. Bauer AW, Kirby WM, Sherris JC, Turck M. 1966. Antibiotic susceptibility testing by a standardized single disk method. Am J Clin Pathol 45:493–496.

19. Baltrus DA, Nishimura MT, Romanchuk A, Chang JH, Mukhtar MS, Cherkis K, Roach J, Grant SR, Jones CD, Dangl JL. 2011. Dynamic evolution of pathogenicity revealed by sequencing and comparative genomics of 19 Pseudomonas syringae isolates. PLoS Pathog 7:e1002132.

20. Bagdasarian M, Lurz R, Rückert B, Franklin FCH, Bagdasarian MM, Frey J, Timmis KN. 1981. Specific-purpose plasmid cloning vectors II. Broad host range, high copy number, RSF 1010-derived vectors, and a host-vector system for gene cloning in Pseudomonas. Gene 16:237–247.

21. Lässig M, Mustonen V, Walczak AM. 2017. Predicting evolution. Nat Ecol Evol 1:77.

22. Heuer H, Smalla K. 2012. Plasmids foster diversification and adaptation of bacterial populations in soil. FEMS Microbiol Rev 36:1083–1104.

23. Zwanzig M, Harrison E, Brockhurst MA, Hall JPJ, Berendonk TU, Berger U. 2019. Mobile Compensatory Mutations Promote Plasmid Survival. mSystems 4.

24. Hall JPJ, Wright RCT, Guymer D, Harrison E, Brockhurst MA. 2019. Extremely fast amelioration of plasmid fitness costs by multiple functionally diverse pathways. Microbiology https://doi.org/10.1099/mic.0.000862.

25. Hooper DC, Jacoby GA. 2016. Topoisomerase Inhibitors: Fluoroquinolone Mechanisms of Action and Resistance. Cold Spring Harb Perspect Med 6.

26. Buckner MMC, Ciusa ML, Piddock LJV. 2018. Strategies to combat antimicrobial resistance: anti-plasmid and plasmid curing. FEMS Microbiol Rev 42:781–804.

27. Winsor GL, Griffiths EJ, Lo R, Dhillon BK, Shay JA, Brinkman FSL. 2016. Enhanced annotations and features for comparing thousands of Pseudomonas genomes in the Pseudomonas genome database. Nucleic Acids Res 44:D646–53.

28. Piddock LJV, Jin YF, Webber MA, Everett MJ. 2002. Novel ciprofloxacin-resistant, nalidixic acid-susceptible mutant of Staphylococcus aureus. Antimicrob Agents Chemother 46:2276–2278.

29. Köhler T, Michea-Hamzehpour M, Plesiat P, Kahr AL, Pechere JC. 1997. Differential selection of multidrug efflux systems by quinolones in Pseudomonas aeruginosa. Antimicrob Agents Chemother 41:2540–2543.

30. Baltrus DA, Dougherty K, Diaz B, Murillo R. 2018. Evolutionary Plasticity of AmrZ Regulation in Pseudomonas. mSphere 3.

31. Orr HA. 2005. The genetic theory of adaptation: a brief history. Nat Rev Genet 6:119–127.

32. Orr HA. 2010. The population genetics of beneficial mutations. Philos Trans R Soc Lond B Biol Sci 365:1195–1201.

33. San Millan A, Toll-Riera M, Qi Q, Betts A, Hopkinson RJ, McCullagh J, MacLean RC. 2018. Integrative analysis of fitness and metabolic effects of plasmids in Pseudomonas aeruginosa PAO1. ISME J 12:3014–3024.

34. Chiotti KE, Kvitek DJ, Schmidt KH, Koniges G, Schwartz K, Donckels EA, Rosenzweig F, Sherlock G. 2014. The Valley-of-Death: reciprocal sign epistasis constrains adaptive trajectories in a constant, nutrient limiting environment. Genomics 104:431–437.

35. Wu NC, Dai L, Olson CA, Lloyd-Smith JO, Sun R. 2016. Adaptation in protein fitness landscapes is facilitated by indirect paths. Elife 5.

36. Molina L, Udaondo Z, Duque E, Fernández M, Molina-Santiago C, Roca A, Porcel M, de la Torre J, Segura A, Plesiat P, Jeannot K, Ramos J-L. 2014. Antibiotic resistance determinants in a Pseudomonas putida strain isolated from a hospital. PLoS One 9:e81604.

37. Gomila M, Peña A, Mulet M, Lalucat J, García-Valdés E. 2015. Phylogenomics and systematics in Pseudomonas. Front Microbiol 6:214.

38. Thomas WJ, Thireault CA, Kimbrel JA, Chang JH. 2009. Recombineering and stable integration of the Pseudomonas syringae pv. syringaeâ61 hrp/hrc cluster into the genome of the soil bacterium Pseudomonas fluorescens Pf0-1. The Plant Journal.

39. Baltrus DA, Medlen J, Clark M. 2019. Identifying transposon insertions in bacterial genomes through nanopore sequencing. Cold Spring Harbor Laboratory.

40. Silva-Rocha R, de Lorenzo V. 2011. A composite feed-forward loop I4-FFL involving IHF and Crc stabilizes expression of the XylR regulator of Pseudomonas putida mt-2 from growth phase perturbations. Mol Biosyst 7:2982–2990.

